# Alteration of DNMT1/DNMT3A by eribulin elicits global DNA methylation changes with potential therapeutic implications for triple-negative breast cancer

**DOI:** 10.1101/2023.06.09.544426

**Authors:** Meisam Bagheri, Min Kyung Lee, Kristen E. Muller, Todd W. Miller, Diwakar R. Pattabiraman, Brock C. Christensen

**Affiliations:** Department of Molecular and Systems Biology, Geisel School of Medicine at Dartmouth, Lebanon, NH 03766; Department of Epidemiology, Geisel School of Medicine at Dartmouth, Lebanon, NH, 03756; Dartmouth Cancer Center, Lebanon, NH, 03756; Department of Community and Family Medicine, Geisel School of Medicine at Dartmouth, Lebanon, NH, 03756; Department of Pathology, Dartmouth-Hitchcock Medical Center, Lebanon NH 03756, USA

**Keywords:** Epithelial-to-mesenchymal transition, mesenchymal-to-epithelial transition, DNA methylation, DNMT1/DNMT3A

## Abstract

Triple-negative breast cancer (TNBC) is an aggressive disease subtype with limited treatment options. Eribulin is a chemotherapeutic approved for the treatment of advanced breast cancer that has been shown to elicit epigenetic changes. We investigated the effect of eribulin treatment on genome-scale DNA methylation patterns in TNBC cells. Following repeated treatment, The results showed that eribulin-induced changes in DNA methylation patterns evident in persister cells. Eribulin also affected the binding of transcription factors to genomic ZEB1 binding sites and regulated several cellular pathways, including ERBB and VEGF signaling and cell adhesion. Eribulin also altered the expression of epigenetic modifiers including DNMT1, TET1, and DNMT3A/B in persister cells. Data from primary human TNBC tumors supported these findings: DNMT1 and DNMT3A levels were altered by eribulin treatment in human primary TNBC tumors. Our results suggest that eribulin modulates DNA methylation patterns in TNBC cells by altering the expression of epigenetic modifiers. These findings have clinical implications for using eribulin as a therapeutic agent.

## Introduction

Triple-negative breast cancer (TNBC) is characterized by the absence of expression of estrogen receptor (ER), progesterone receptor (PR), and human epidermal growth factor receptor 2 (HER2). TNBC accounts for approximately 15% of all breast cancer cases and is associated with poorer prognosis compared to other subtypes due to its aggressive nature and lack of targeted therapy (1, 2). The current standard of care for most TNBC cases includes a combination of conventional chemotherapy, radiation therapy, immunotherapy, and surgery (2, 3).

DNA methylation is an epigenetic mark known for its functions in regulating transcription and occurs on cytosine residues that are linked to a guanine base (CpGs). DNA methylation is controlled mainly by two enzyme families: DNA methyltransferases (DMNT1, DNMT3A, DNMT3B) and tet methylcytosine dioxygenases (TET1, TET2, TET3; also known as ten-eleven translocation enzymes). DMNTs add methyl groups to cytosine residues, and TET enzymes oxidize those methyl groups as a part of an active demethylation pathway (4). Dysregulation of these enzymes has been implicated in developing and progressing cancers including TNBC(5, 6). Altered DNA methylation patterns have been shown to contribute to oncogene activation and the silencing of tumor suppressors, driving the aggressive behavior of TNBC (7, 8). Overexpression of DNMT1 and DNMT3B has been observed in TNBC, leading to hypermethylation, and silencing of tumor suppressors, whereas the level of TET1 is downregulated in TNBC (9-11). Moreover, DNA methylation patterns can potentially serve as biomarkers for early detection, prognosis, and treatment response in TNBC (8). Therapeutic targeting of such altered enzymes may restore pre-cancer DNA methylation patterns and improve TNBC outcomes. Indeed, DNMT inhibitors such as azacytidine and decitabine that are currently approved for the treatment of patients with hematological malignancies have been suggested to improve outcomes for TNBC patients (12,13).

Eribulin is a chemotherapeutic microtubule dynamics inhibitor approved by the US Food and Drug Administration (FDA) for the treatment of patients with metastatic breast cancers, including TNBC (14). Clinical trials demonstrated that eribulin is associated with improved overall survival (14.4 versus 9.4 months) in TNBC patients (15). Preclinical work showed that eribulin induces a mesenchymal-to-epithelial transition, and treatment of cancer cells with eribulin leads to the formation of epithelial-like persisters compared to both parental cells and those treated with paclitaxel (16). Epithelial-to-mesenchymal transition (EMT) and its reverse process, mesenchymal-to-epithelial transition (MET), are recognized as crucial events in cancer progression and metastasis. EMT enables cancer cells to acquire invasive and migratory properties, facilitating their escape from the primary tumor site and dissemination to distant organs. Conversely, MET allows disseminated cancer cells to undergo a phenotypic transition, enabling them to establish secondary tumor colonies in new microenvironments. The dynamic interplay between EMT and MET contributes to tumor heterogeneity, plasticity, treatment resistance, and poor clinical outcomes for cancer patients (17,18). Previously, we observed significant changes in transcriptional programs upon eribulin selection. This led us to hypothesize that these changes were a result of modifications to the chromatin state in cancer cells. Specifically, we found a reduction in the levels of the H3K4me3 activation mark and the H3K27me3 repressive mark in ERI-R derivatives, while PAC-R derivatives showed no such alterations. Considering the established association between alterations in transcriptional programs and DNA methylation patterns with epithelial-to-mesenchymal transition (16,19), we posit that eribulin’s potential anti-tumor effects in triple-negative breast cancer (TNBC) may be attributed to its capacity to modulate DNA methylation patterns and regulate genes implicated in cancer progression. Targeting EMT and MET pathways holds promise for developing novel therapeutic strategies aimed at inhibiting cancer metastasis and improving patient outcomes. By unraveling the impact of eribulin on DNA methylation and its associated enzymes, we can gain valuable insights into the therapeutic potential of this drug as a targeted therapy for TNBC.

## Results

### Eribulin treatment alters DNA methylation patterns in TNBC cells

To investigate the effect of eribulin on DNA methylation, we treated MDA-MB 231 TNBC cells with two rounds of eribulin to generate persister-epithelial cancer cells. As a comparison, we also treated cells with two rounds of paclitaxel, another microtubule inhibitor (**Figure 1A**). We then measured epigenome-wide DNA methylation patterns at each time point. Principal components analysis (PCA) showed that DNA methylation profiles of eribulin-treated cells were distinct from those of parental cells and paclitaxel-treated cells (**Figure 1B** and **Supplementary Figure 1A**). Supporting the PCA results, unsupervised hierarchical clustering of the top 5% most variable CpGs (34,525 CpGs) indicated methylation profiles of eribulin-treated cells to be distinct from profiles of paclitaxel-treated and parental cells (**Figure 1C**). To quantify the cumulative burden of eribulin and paclitaxel treatments on DNA methylation, we calculated the methylation dysregulation index (MDI, a summary measure of cumulative departure of DNA methylation levels) compared to untreated TNBC cells. The MDI of eribulin-treated cells (after ERI1/2 vs PAC1/2) was higher than that of paclitaxel-treated cells (two-sided Wilcoxon test, p-value = 0.03), and MDI increased with the second round of eribulin treatment compared to the first round (ERI1 vs ERI2) (**Figure 1D**).

**Figure 1.**
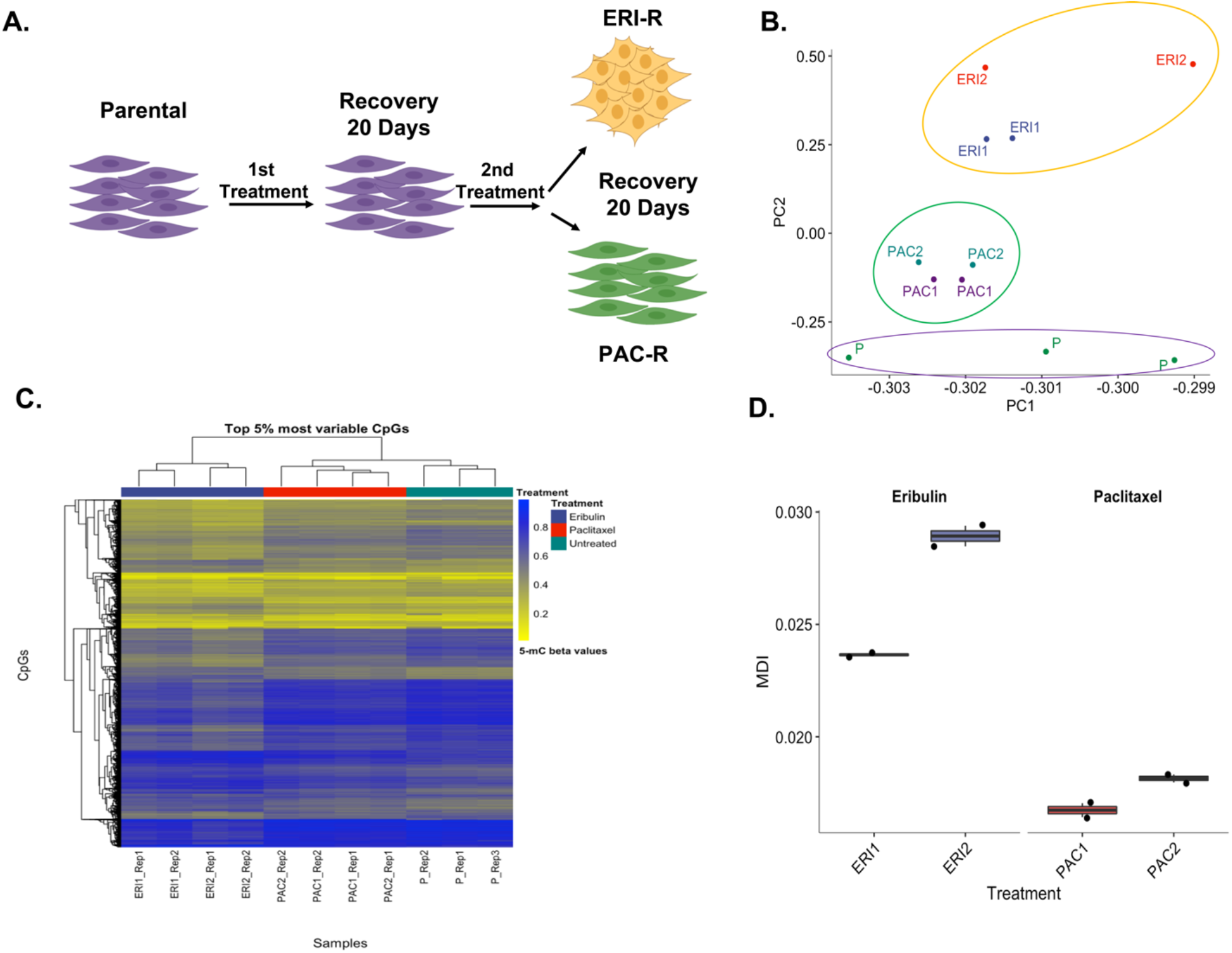
Eribulin treatment alters DNA methylation patterns in TNBC cells. (A) Schematic representation of multi-round drug treatment procedure that resulted in the generation of resistant clones (B) Principal Component Analysis (PCA) of most variable genes between MDA MB-231 parental, ERI1, ERI2, PAC1 and PAC2 from DNA methylation array. (C) (D)

To determine which differentially methylated CpGs (dmCpGs) were associated with eribulin and paclitaxel treatments, we conducted epigenome-wide association tests. After the first round of eribulin, 47,758 dmCpGs were detected (26,383 hypomethylated; 21,375 hypermethylated) (**Figure 2A**). The number of dmCpGs increased to 101,949 after the second round of eribulin (64,351 hypomethylated CpGs; 37,598 hypermethylated CpGs) (**Figure 2B**). There were more hypomethylated CpGs after each eribulin treatment than hypermethylated CpGs (first round: 55% of dmCpGs; second round: 63% of dmCpGs; **Figure 2C**). The first and second rounds of eribulin resulted in the same direction of change in methylation levels at nearly all methylated CpGs (**Figure 2D**). 24,024 hypomethylated CpGs (91% of single-treatment sites) and 17,003 hypermethylated CpGs (80% of single-treatment sites) were shared between the first and second eribulin treatments. Furthermore, the dmCpGs associated with the additional eribulin treatment were located within very similar sets of genes as the dmCpGs related to the initial eribulin treatment, as thousands of genes had an increase in the proportion of dmCpGs per gene (**Figure 2E**). The biological processes associated with hypermethylated and hypomethylated CpGs across both eribulin hits were similar. dmCpGs associated with eribulin treatments were associated with differentiation, organismal processes, and system development (**Supplementary Figure 2**).

**Figure 2.**
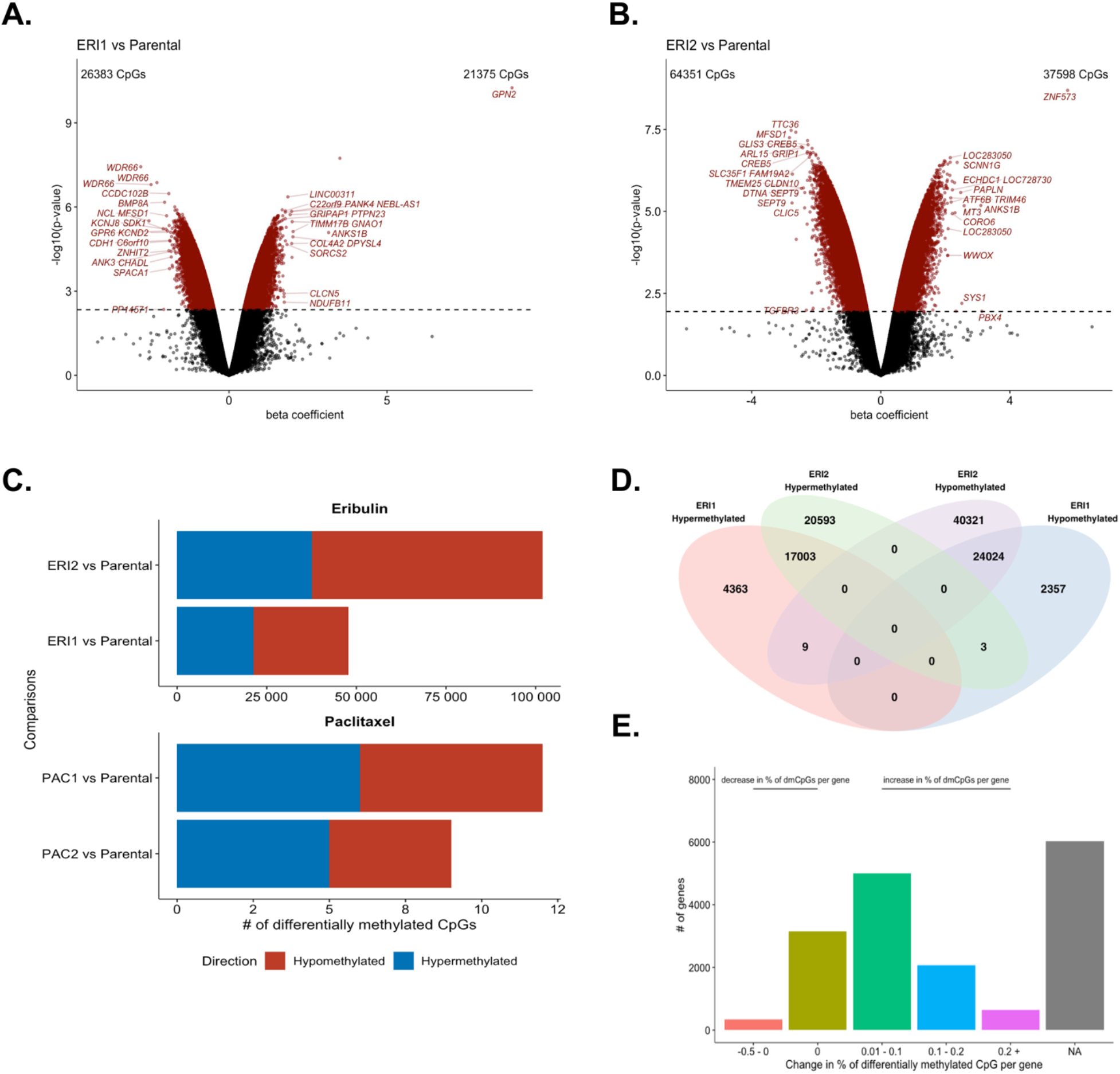
Additional eribulin treatment increases alterations in DNA methylation patterns. (A) Volcano plot of dmCpGs associated with first eribulin treatment compared to the methylation status of parental cells determined by an epigenome-wide association study. (B) Volcano plots of dmCpGs associated with second eribulin treatment compared to the methylation status of parental cells determined by an epigenome-wide association study. Colored in red are considered to be differentially methylated at q-value < 0.05. (C) Comparison of number of dmCpGs based on treatment type and number of treatments. Colored in red are number of hypomethylated CpGs compared to the methylation status of parental cells. Colored in blue are number of hypermethylated CpGs compared to methylation status of parental cells. (D) Venn diagram of dmCpGs associated with the first treatment of eribulin and associated with the second treatment of eribulin. (E) Number of genes that have either decreased, unchanged, or increased proportion of CpGs that are differentially methylated with the second treatment of eribulin as compared to the proportion of CpGs that are differentially methylated with the first treatment of eribulin.

While tens of thousands of CpGs were identified to be differentially methylated in eribulin-treated TNBC cells compared to untreated cells, we observed very few dmCpGs following paclitaxel treatment. Only 12 CpGs were differentially methylated (6 hypomethylated CpGs; 6 hypermethylated CpGs) with the first paclitaxel treatment compared to baseline (**Supplementary Figure 3A**). Nine CpGs were differentially methylated (4 hypomethylated CpGs; 5 hypermethylated CpGs) after the second round of paclitaxel treatment (**Supplementary Figure 3B**).

To identify patterns of differential methylation levels at specific genomic loci, we tested dmCpGs induced by drug treatment for enrichment at various genomic contexts, first in context in relation to the gene structure and second, in context in relation to the location of the CpG. Both hypermethylated and hypomethylated CpGs associated with eribulin treatment were significantly enriched for tracking to enhancer regions (**Figure 3A**). Hypermethylated CpGs were enriched in open chromatin regions. On the contrary, hypomethylated CpGs were enriched in transcription factor binding sites (TFBS). While hypermethylated CpGs were enriched in shore/shelf regions and open sea (inter CpG island regions, or regions of less dense CpGs clustered together) regions, hypomethylated CpGs were enriched only in open sea regions (**Figure 3B**).

**Figure 3.**
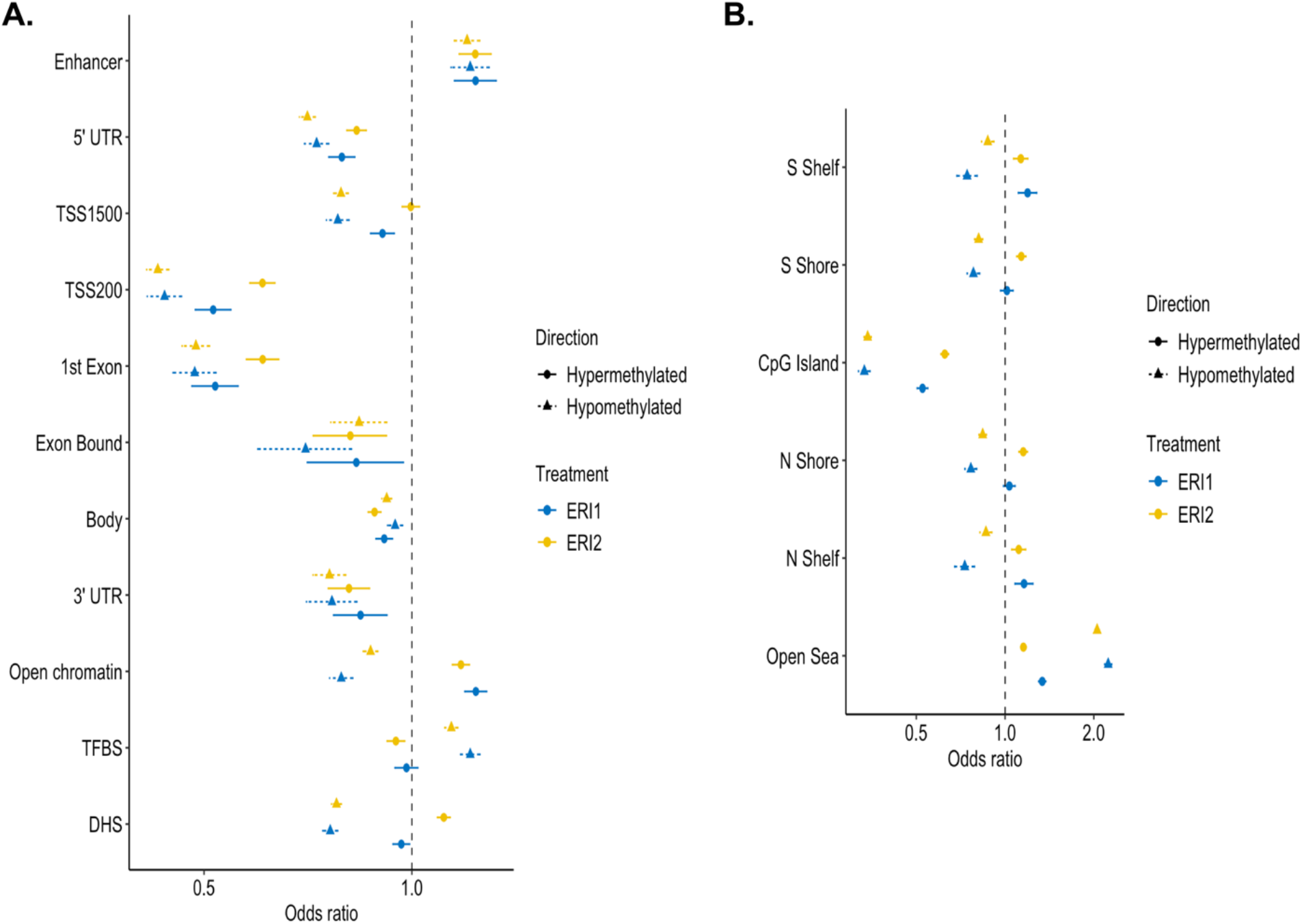
Eribulin treatment associated with differentially methylated CpGs are enriched. Enrichment of dmCpGs at different genomic contexts (A) in relation to the gene structure and (B) in relation to the CpG islands. Odds ratios were calculated by Fisher’s exact test. Blue points and confidence intervals indicate enrichment from dmCpGs associated with the first eribulin treatment. Yellow points and confidence intervals indicate enrichment from dmCpGs associated with the second eribulin treatment. Circular points indicate CpGs with increased methylation levels in treated cells. Triangular points indicate CpGs with decreased methylation levels in treated cells.

We aimed to investigate signaling pathways significantly impacted by eribulin. We identified 91 pathways associated with genes that had that were significantly hypermethylated CpGs, and 57 pathways associated with genes that had significantly hypomethylated CpGs (q-value < 0.05; **Table S1, S2**). Specifically, our analysis showed that eribulin treatment led to differential methylation of CpGs in genes involved in the ERBB signaling pathway, including *ERBB2* that encodes HER2 (**Figure 4A; Supplementary Figure 4A**), and the vascular endothelial growth factor (VEGF) signaling pathway (**Figure 4B, Table S1, S2**). Furthermore, eribulin caused altered methylation levels of the CpGs in *VEGF* genes and altered ZEB1 interactions with NF-kappa B1/2 transcription factors locus (**Figure 4D – 4F, Table S3, S4**). To explore these findings further, we conducted a multiplexed cytokine array analysis and observed notable alterations in cytokine expression levels in eribulin-treated cells compared to the parental cells. Specifically, CXCL10, GM-CSF, and VEGF exhibited significant decreases, corroborating the notion of increased DNA methylation within these pathways (**Figure 4C**). In addition, we observed differential methylation at CpGs of genes involved in cell adhesion, focal adhesion, and gap junction pathways after both first and second treatment of eribulin (**Supplementary Figure 4B – 4C**). These results suggest that eribulin may suppress the ERBB and VEGF signaling pathways while promoting cell adhesion and intercellular communication through alterations in DNA methylation patterns. These findings provide insights into the mechanisms underlying eribulin-induced cellular changes and suggest potential therapeutic strategies for sensitizing resistant cancer cells.

**Figure 4.**
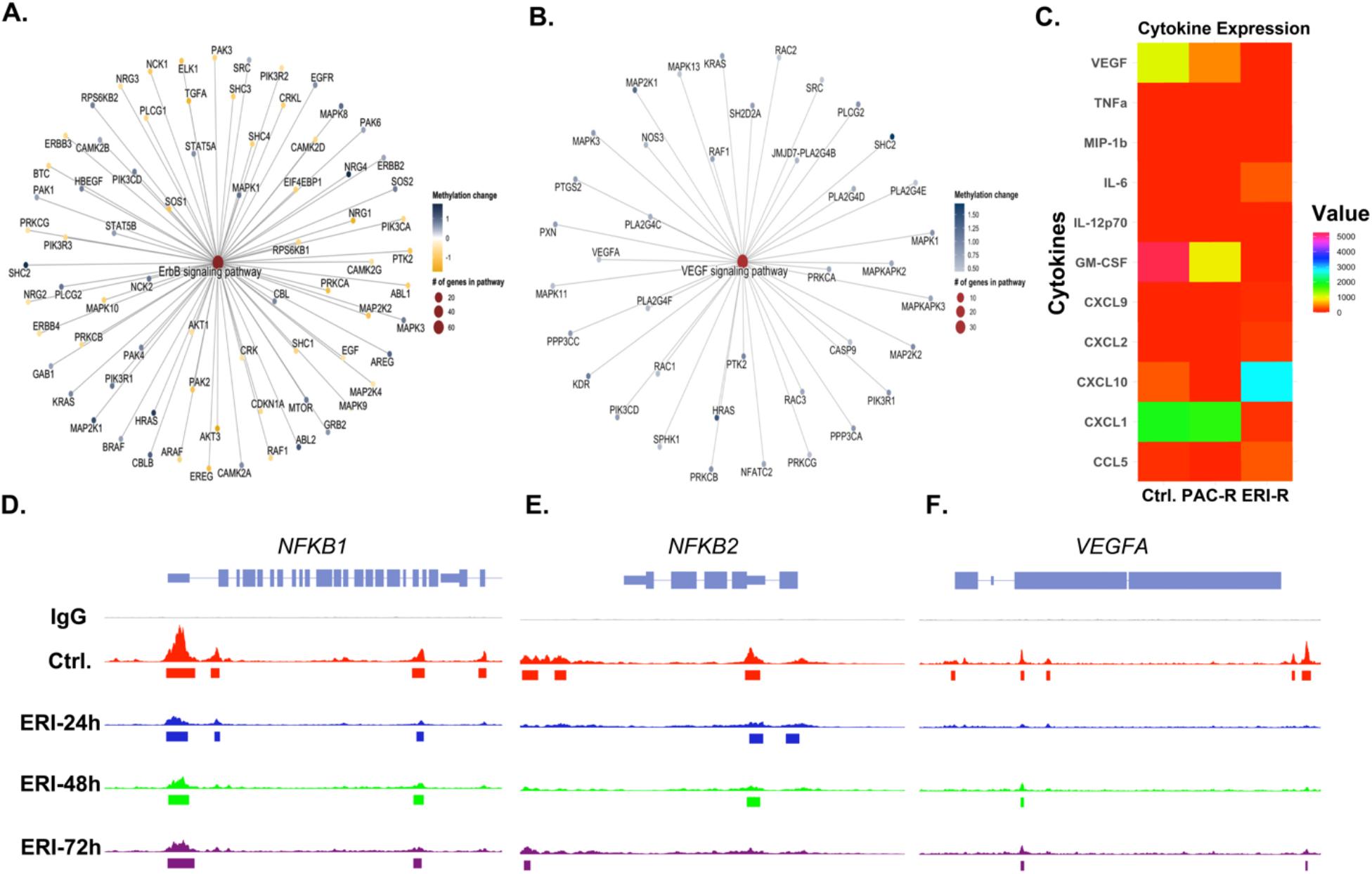
Genes with differentially methylated CpGs are associated with key biological processes in the mesenchymal-to-epithelial transitions. (A) ERBB signaling pathway was significant associated with dmCpGs from the second eribulin treatment. (B) VEGF signaling pathway were associated with hypermethylated CpGs from the second eribulin treatment. (C) Heatmap depicting the levels of cytokines and chemokines determined through a multiplex cytokine assay. ZEB1 binding levels at (D) *NKFB1*, (E) *NKFB2*, and (F) *VEGFA* in untreated and eribulin treated cells.

### Differential methylation patterns are detected in Mesenchymal to Epithelial Transition-associated genes

Among 21 known mesenchymal-to-epithelial transition (MET)-associated genes (*CDH1, CLDN1, EPCAM, ITGB4, KRT8, KRT14, OCLN, CDH2, FN1, ITGB1, MMP19, MMP2, VIM, SNAI1, SNAI2, TWIST1, ZEB1, ZEB2, OVOL1, OVOL2, GHRL*), 40 dmCpGs in MET-associated genes were associated with the first eribulin treatment, and 94 dmCpGs in MET-associated genes were associated with the second eribulin treatment. Generally, CpGs were hypomethylated in eribulin-treated cells across EMT-associated transcription factor genes (*SNAI1, SNAI2, TWIST1, ZEB1, ZEB2*; **Figure 5A**). Like other dmCpGs, the number of dmCpGs increased with additional eribulin treatment in 80% of the EMT-associated genes (**Supplementary Figure 5**). A large proportion of dmCpGs were located in the gene body and DNase-hypersensitive sites of EMT-associated genes, and moderate proportions of dmCpGs were located in promoter regions (within 1500 bp of transcription start sites) of EMT-associated genes (**Figure 5B**).

**Figure 5.**
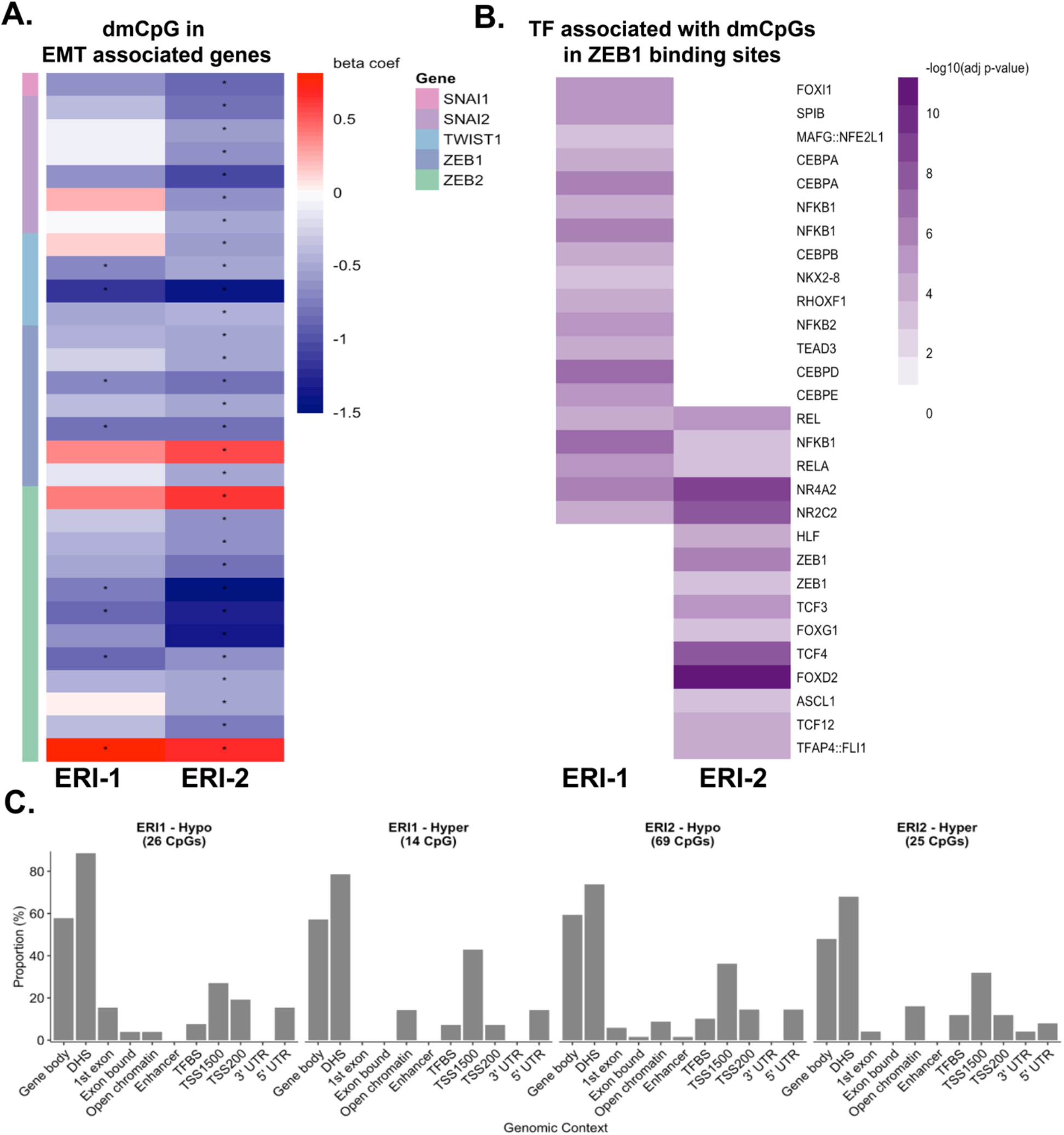
Differential methylation patterns are detected in Mesenchymal to Epithelial Transition-associated genes. (A) Heatmap of the direction of methylation level change (beta coefficient from epigenome-wide association study) in CpGs located in MET-associated genes. Tiles with * indicate a significant difference in methylation levels compared to untreated cells. The vertical tracking bar indicates which MET-associated gene the CpG is located in. (B) Proportion of dmCpGs in different genomic regions of MET-associated genes by direction of methylation change and number of eribulin treatment. (C) Heatmap of the level of significance for transcription factor motifs associated with dmCpGs in ZEB1 binding sites.

We found the differentially methylated CpG sites (dmCpGs) to be enriched in enhancers, transcription factor binding sites, and open chromatin regions (**Figure 3A**). In addition, as our previous findings showed that eribulin can alter the ZEB1-SWI/SNF interaction and induce mesenchymal-to-epithelial transition (MET) (16). So, we aimed to identify if differentially methylation would affect any other transcription factors at ZEB1 binding sites. To accomplish this, we performed transcription factor motif enrichment testing at ZEB1 binding sites (identified from our CUT&RUN experiment) containing dmCpGs. Our analysis revealed that after the first eribulin treatment, there were 19 significantly enriched transcription factor motifs for dmCpG-containing ZEB1 binding sites. In contrast, after the second round of eribulin, there were 15 significantly enriched transcription factor motifs (**Figure 5C**). Interestingly, in TNBC cells treated with the first round of eribulin, motifs for CEBP proteins (CEBPA, CEBPB, CEBPD, CEBPE) and NFkB1/2 were significantly enriched. Conversely, in TNBC cells treated with the second eribulin hit, motifs for TCF proteins (TCF12, TCF3, TCF4) and ZEB1 were enriched. These findings suggest that eribulin treatment may affect the binding of specific transcription factors to ZEB1 binding sites or the binding of ZEB1 to different regions containing dmCpGs. These results provide new insights into the molecular mechanisms underlying eribulin-induced MET and highlight the importance of understanding the regulatory networks involved.

### Eribulin treatment in primary human triple-negative breast cancers alters the expression of DNA methylation regulatory enzymes

To explore the mechanism underlying eribulin-induced modification of DNA methylation in MDA-MB-231 cells, we employed RT-qPCR analysis to measure the expression of enzymes regulating DNA methylation. Our results revealed a significant increase in the expression of *TET1, DNMT3A*, and *DNMT3B* in eribulin-treated cells compared to parental and paclitaxel-treated cells. In contrast, the expression of *TET2, DNMT1*, and *DNMT3I* remained unchanged or was downregulated (**Supplementary Figure 6A**). Our findings were confirmed by immunoblot analysis (**Figure 6A**).

**Figure 6.**
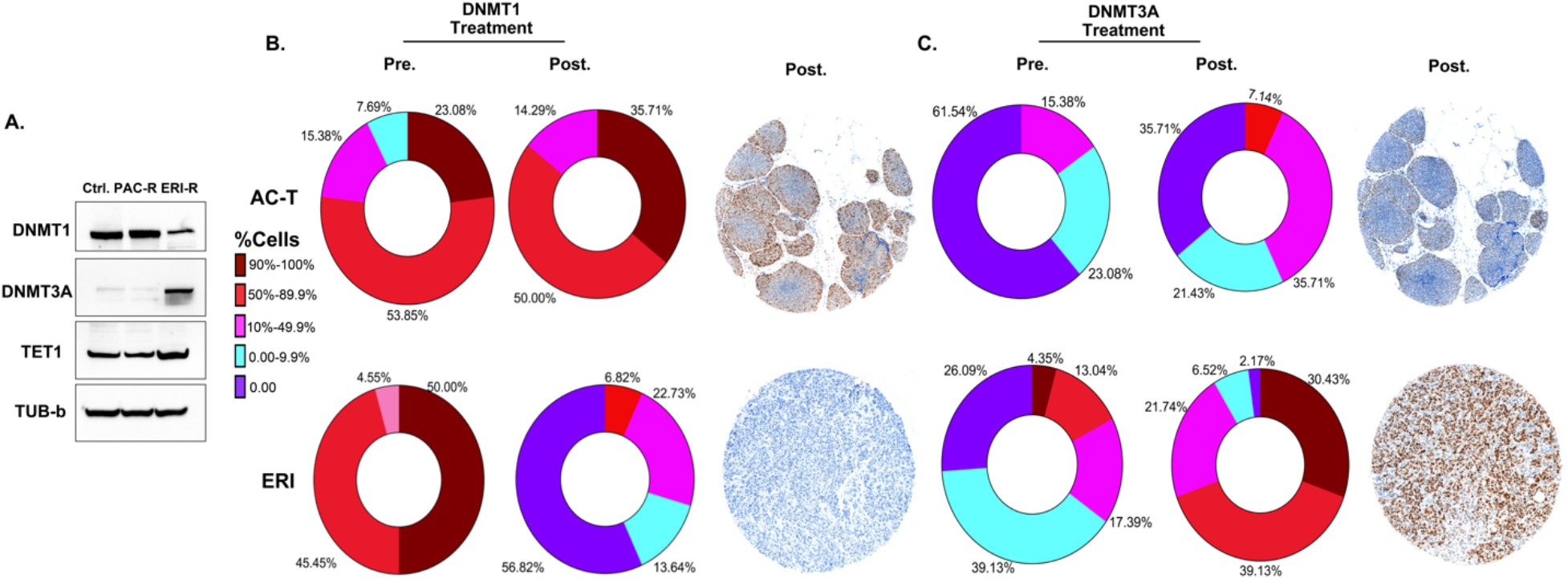
Eribulin treatment in primary human triple-negative breast cancers alters the expression of DNA methylation regulatory enzymes. (A) Immunoblotting was performed to assess the protein levels of DNA methylation markers in the MDA MB-231 parental line and its resistant counterparts. (B, C) Representative images and quantification of DNMT1 and DNMT3A were obtained through immunohistochemistry (IHC) analysis of specimens treated with AC-T and ERI. Statistical analysis was conducted using the Wilcoxon-Mann-Whitney test.

To validate the clinical relevance of our results, we conducted immunohistochemical staining (IHC) for *TET1, DNMT1*, and *DNMT3A* on TNBC diagnostic core biopsy specimens (pre-treatment) and surgical specimens (post-treatment) obtained from patients who received neoadjuvant chemotherapy. The specimens from 55 patients who received 4 cycles of neoadjuvant eribulin (SOLTI1007 NeoEribulin clinical trial)(20) were compared with specimens from 15 patients who received standard-of-care neoadjuvant treatment with 4 cycles of adriamycin/cyclophosphamide followed by 4 cycles of paclitaxel (AC-T). Results confirmed our *in vitro* findings, demonstrating that the level of DNMT1 remained unchanged in AC-T-treated tumors (**Figure 6B**). However, in eribulin-treated specimens, DNMT1 levels were significantly reduced in approximately 70% of tumors compared to baseline, with 56.82% of specimens showing complete loss of expression and 13.64% exhibiting expression in ≤10% of malignant cells. The levels of DNMT3A did not show significant changes in AC-T-treated specimens; However, a notable increase in DNMT3A expression was observed in eribulin-treated specimens. Specifically, among post-eribulin specimens, 30.43% of tumors displayed 90-100% and 39.13% demonstrated 50-89.9% DNMT3A expression. In comparison, pre-treated samples exhibited 4.35% and 13.04% DNMT3A expression in 90-100% and 50-89.9% of malignant cells, respectively (**Figure 6C**). Compared to our *in vitro* data, levels of TET1 did not show changes in any of the specimens (**Supplementary Figure 6B**).

## Discussion

These data collectively suggest that eribulin treatment affects the global DNA methylation of breast cancer cells by upregulating DNMT3A/B and downregulating DNMT1 expression. Our previous study demonstrated that eribulin treatment of TNBC cells induces significant changes in chromatin and gene expression patterns (16). Given the critical role of DNA methylation in gene expression regulation and genomic stability maintenance, we investigated the DNA methylation patterns of eribulin-treated cells to understand further the molecular mechanisms underlying these changes (21). Our findings demonstrate that eribulin treatment induces distinct changes in DNA methylation patterns in MDA-MB 231 cells compared to parental and paclitaxel-treated cells. These changes are more pronounced in persisters that arise following repeated eribulin treatment. These results are consistent with previous studies that have reported chemotherapy-induced epigenetic changes, including DNA methylation alterations (5,21). Interestingly, our study revealed that eribulin treatment leads to a higher degree of methylation dysregulation compared to paclitaxel treatment, which may contribute to the observed mesenchymal-to-epithelial transition and formation of epithelial-like persisters (16,19).

Moreover, the number of differentially methylated CpGs increased with additional eribulin treatments, with a higher proportion of hypomethylated CpGs compared to hypermethylated CpGs. This finding supports the concept of epigenetic plasticity (8), which refers to the ability of cancer cells to adapt to environmental changes by altering their epigenetic landscape. On the other hand, very few CpGs were differentially methylated with paclitaxel treatment, which is consistent with previous studies demonstrating that paclitaxel treatment does not significantly alter DNA methylation patterns in breast cancer cells (22,23). These results suggest that eribulin treatment induces significant changes in DNA methylation patterns that may contribute to the acquisition of drug resistance in cancer cells. Moreover, they highlight the potential of epigenetic therapies as a complementary approach to conventional chemotherapy to treat TNBC (24,25). Further studies are needed to investigate the functional consequences of eribulin-induced DNA methylation changes and their role in drug resistance acquisition in cancer cells.

In this study, we aimed to identify patterns of differential DNA methylation associated with eribulin treatment in triple-negative breast cancer (TNBC) cells. Our results revealed both hypermethylated and hypomethylated CpGs associated with eribulin treatment enriched in enhancers, with hypermethylated CpGs also enriched in open chromatin regions and hypomethylated CpGs enriched in transcription factor binding sites. Transcription factor motif enrichment tests revealed that eribulin treatment might affect the binding of specific transcription factors to ZEB1 binding sites or the binding of ZEB1 to different regions containing dmCpGs, providing new insights into the molecular mechanisms underlying eribulin-induced mesenchymal-to-epithelial transition (MET) (26,27). Additionally, our findings suggest that eribulin treatment can regulate multiple cellular pathways, including the ERBB signaling pathway, VEGF signaling pathway, cell adhesion, focal adhesion, and gap junction pathways, by altering DNA methylation patterns (28). Specifically, eribulin treatment led to hypermethylation of genes involved in the ERBB and VEGF signaling pathways, critical pathways for cancer cell survival, proliferation, and metastasis (29), and hypomethylation of genes involved in maintaining normal cellular functions and tissue architecture. Moreover, our results showed that eribulin treatment altered the interactions between ZEB1 and NFkB1/2 transcription factors, highlighting its potential role in regulating EMT and providing new insights into the molecular mechanisms underlying eribulin-induced MET (30).

Our study revealed that eribulin persister cells exhibited downregulation of DNMT1 expression, which is responsible for maintaining DNA methylation patterns, and upregulation of TET1 and DNMT3A/B expression involved in active DNA demethylation and de novo DNA methylation, respectively (31). Our clinical study further supported these findings by demonstrating that eribulin-treated specimens had higher levels of DNMT3A and lower levels of DNMT1 compared to pre-treatment specimens. These changes in gene expression suggest that eribulin treatment may alter DNA methylation patterns, leading to potential effects on gene expression and ultimately contributing to the drug’s therapeutic efficacy.

The implications of altered DNA methylation patterns in cancer have been well-documented (32, 33). In breast cancer, aberrant DNA methylation patterns have been linked to tumor aggressiveness, and targeting DNA methylation pathways has been explored as a therapeutic strategy (34). Our study adds to this body of literature by providing evidence that eribulin treatment may affect global DNA methylation patterns in TNBC. Our clinical data also support our in vitro findings, as we observed changes in the expression of DNMT1 and DNMT3A in TNBC specimens obtained from patients treated with eribulin compared to those treated with standard-of-care chemotherapy. These findings are consistent with previous studies that have reported alterations in DNA methylation patterns in response to chemotherapy (35,36).

In conclusion, our study sheds light on the mechanism of action of eribulin in breast cancer and suggests that eribulin treatment alters DNA methylation patterns in primary human TNBC by upregulating DNMT3A/B and downregulating DNMT1 expression. These findings have important clinical implications for the potential use of eribulin as a therapeutic agent for breast cancer. Further studies are needed to fully elucidate the molecular mechanisms underlying the effects of eribulin on DNA methylation and to determine whether these changes contribute to eribulin’s therapeutic efficacy in breast cancer.

## Acknowledgments

We thank Dr. Owen Wilkins at the Center for Quantitative Biology at Dartmouth for providing code for statistical analysis. We thank the Dartmouth Cancer Center Genomics and Molecular Biology Shared Resource and Pathology Shared Resource. Funding and resources for the shared resources were supported in part by a core grant (P30CA023108; Dartmouth Cancer Center). Research reported in this publication was supported through the Geisel School of Medicine at Dartmouth’s Center for Quantitative Biology through a grant from the NIGMS Award P20GM130454, an NIH S10 (S10OD025235) grant, funding from The Elmer R. Pfefferkorn & Allan U. Munck Education and Research Fund at the Geisel School of Medicine at Dartmouth (D.R.P), and a Sponsored Research Agreement with Eisai Inc. (D.R.P). This work was supported by funding from the NIH R01GM122846 (S.A.G), R00CA201574 (D.R.P.), METAvivor (D.R.P), and R01CA267691 (D.R.P. and T.W.M.). R01CA216265 to BCC.

## Author contributions

### Conception and design

M.B., M.K.L.

### Development of methodology

M.B., M.K.L, K.E.M., T.W.M., B.C.C.

### Acquisition of data

M.B., M.K.L.

### Analysis and interpretation of data

M.B., M.K.L, K.E.M., T.W.M., B.C.C.

### Writing, review, and/or editing of the manuscript

M.B., M.K.L, K.E.M., T.W.M., B.C.C., D.R.P.

### Study supervision

T.W.M., B.C.C., D.R.P.

### Declaration of Conflicts of Interest

This study was supported in part by a Sponsored Research Agreement with Eisai Inc.

### Data and materials availability

## Methods

### Cell lines and culture conditions

The MDA-MB-231 cell line was a gift from Dr. Bob Weinberg (Whitehead Institute for Biomedical Research). MDA-MB-231 and SUM-159 were cultured in DMEM supplemented with 10% FBS, per ATCC guidelines.

### Drug Treatment

Stock solutions for eribulin (10 mM) were prepared using double distilled water, and paclitaxel was prepared using DMSO, further diluted in the base diluent to the appropriate concentrations. The half-maximal inhibitory concentration (IC50) was determined at different concentrations from 0.5nM to 100nM using a 2-fold dilution series for MDA-MB-231 cells.

### CUT&RUN

CUT&RUN was performed using CUTANA™ Kit (EpiCypher #14-1048) with minor modifications according to the manufacturer’s recommendation. Briefly, 5×105 cells were washed and bound to concanavalin A-coated magnetic beads. The cells were then permeabilized with Wash Buffer (20 mM HEPES pH 7.5, 150 mM NaCl, 0.5 mM spermidine, and 1x Roche Complete Protease Inhibitor, EDTA-free) containing 0.025% digitonin (Digitonin Buffer) and 2 mM EDTA and incubated with primary antibody (anti-ZEB1 or IgG isotype control) overnight at 4°C. The cell-bead slurry was washed twice with Digitonin Buffer and incubated with 1x Protein-A/G-MNase (pAG-MNase) in Digitonin Buffer for 10 minutes at room temperature. The slurry was washed twice with Digitonin Buffer and incubated in Digitonin Buffer containing 2 mM CaCl2 for 2 hours at 4°C to activate pAG-MNase digestion. The digestion was stopped by addition of 2x Stop Buffer (340 mM NaCl, 20 mM EDTA, 4 mM EGTA, 50 μg/mL RNase A, 50 μg/mL glycogen), and the sample was incubated for 10 minutes at 37°C to release chromatin to the supernatant and degrade RNA. The supernatant was recovered, and DNA was isolated using MinElute Reaction Cleanup Kit (Qiagen # 28206). Isolated CUT&RUN DNA fragments were quantified by Qubit and 5-10ng used for library preparation with the NEB Ultra II DNA Kit. Library amplification was performed using the modified PCR cycling conditions described in Step 39 of the Epicypher CUTANA™ (EpiCypher) protocol. Libraries individually barcoded and pooled for sequencing on a NextSeq500 Mid Output flow cell to generate 10 million, 50-bp paired-end reads per sample and analyzed as described in 16.

### Immunoblotting

Protein extraction was performed by pelleting cells and resuspending in cold RIPA buffer (Thermo Fisher Scientific, #89901) supplemented with phosphatase/protease inhibitors (Thermo Fisher Scientific, #78446) and incubating for 1 hour on ice. Then, protein extracts were collected at 20,000 rcf for 10 min in a refrigerated benchtop centrifuge. Protein lysates were quantified using Quick Start™ Bradford Protein Assay kit (BIO-RAD #5000202) and then diluted NuPAGE LDS sample buffer (Life Technologies #NP0007) and NuPAGE™ Sample Reducing Agent (Life Technologies #NP0009) and heated at 72°C before resolving on 4-12% Bis-Tris gradient gels. Gels were either wet or dry transferred using Mini Blot module or iBlot™ 2 Gel Transfer Device, respectively (Thermo Fisher Scientific #B1000 and #IB21001). All primary antibodies were incubated overnight with membranes in TBS blocking buffer supplemented with 0.1% Tween-20 (Sigma-Aldrich # P1379), while secondary antibodies (HRP conjugate) were incubated at room temperature with agitation for 1 hour in the primary blocking buffer. Membranes were developed using SuperSignal™ Western Blot Substrate (Thermo Fisher Scientific, # A45915). (List of antibodies were used: Cell Signaling Technology DNMT1 #5032, DNMT3A #3598 α/β-Tubulin #2148, GeneTex TET1 # 124207).

Tissue samples were collected from [source] and fixed in 10% neutral buffered formalin for 24-48 hours. The fixed tissue samples were then dehydrated in a series of graded ethanol solutions (70%, 80%, 95%, and 100%) for 30 minutes each, cleared in xylene for 30 minutes, and embedded in paraffin wax. Using a microtome, paraffin-embedded tissue blocks were sectioned at 4-5 μm thickness.

### Immunohistochemistry

Tissue sections were deparaffinized in xylene for 10 minutes and rehydrated in a series of graded ethanol solutions (100%, 95%, 80%, and 70%). Antigen retrieval was performed by heating the tissue sections in a citrate buffer (pH 6.0) at 95-100°C for 20 minutes. Endogenous peroxidase activity was blocked by incubating the sections in 3% hydrogen peroxide for 10 minutes, followed by blocking of non-specific binding sites using 10% normal serum (diluted in PBS) for 30 minutes at room temperature.

The primary antibodies (Cell Signaling Technology DNMT1 #5032, DNMT3A #3598, GeneTex TET1 # 124207) were applied to the sections and incubated at 4°C overnight. The following day, sections were washed with PBS three times for 5 minutes each, and a biotinylated secondary antibody was applied for 1 hour at room temperature. The sections were washed with PBS three times for 5 minutes each and then incubated with an avidin-biotin-peroxidase complex (ABC kit, dilution) for 30 minutes at room temperature.

Peroxidase activity was visualized by incubating the sections with 3,3’-diaminobenzidine (DAB) for 5 minutes, followed by counterstaining with hematoxylin for 2 minutes. The sections were then dehydrated in a series of graded ethanol solutions (70%, 80%, 95%, and 100%) for 30 seconds each, cleared in xylene for 5 minutes, and mounted with a coverslip using a mounting medium.

Stained tissue sections were observed under a microscope, and images were captured for analysis. The staining patterns and intensities were analyzed, and the staining was quantified by measuring the optical density using image analysis software. Statistical analysis was performed to determine the significance of the staining differences observed.

### DNA methylation data collection and pre-processing

DNA were converted with bisulfite treatment and hybridized to Illumina HumanMethylation EPIC array. Raw idat files from the methylation array were pre-processed with SeSAMe Bioconductor pipeline for normalization and beta value calculation (cite). Quality control was performed using *ENmix* R package (cite). Low quality probes, SNP-associated probes and cross-reactive probes were removed for downstream analyses (cite Zhou et al). 690,483 CpGs remained after these exclusions.

### Statistical analysis

Methylation beta values were logit2 transformed to M-values for a more normal distribution of the methylation dataset with the *logit2* function in the *minfi* R package. Epigenome wide association studies to identify differentially methylated CpGs associated with the different treatment groups were identified by fitting M-values into linear regression models. Linear regression models were fit by using *lmFit* and *eBayes* functions in *limma* R package. P-values were adjusted for multiple comparisons to q-values using the *qvalue* R package. Differentially methylated CpGs were determined to be significant under the q-value threshold of 0.05.

Enrichment at different genomic contexts for CpGs was tested compared to the 690,483 CpGs using Fisher’s exact tests. Annotations of CpGs for genomic context were provided in the *IlluminaHumanMethylationEPICanno*.*ilm10b5*.*hg38* R package (37). Genomic Regions Enrichment of Annotations Tool (GREAT) was used to assess the functional significance of the differentially methylated CpGs (38). enrichKEGG function in *clusterProfiler* R package was used to determine pathways associated with genes that had associated differentially methylated CpGs(39,40). Over-representation of transcription factor binding site motifs with differentially methylated CpGs within the ZEB1 binding sites were tested against all ZEB1 binding sites determined from ZEB1 CUT&RUN experiments using hypergeometric testing using the *phyper* R package. Transcription factor binding site motifs curated in the JASPAR database, downloaded with *JASPAR2022* and *TFBStools* R packages were used for analyses (41,42). TF motifs with FDR-adjusted p-value < 0.05 were considered to be over-represented.

## Notes

### Competing Interest Statement

The authors have declared no competing interest.

