## Supplementary Figures for "Alteration of DNMT1/DNMT3A by eribulin elicits global DNA methylation changes with potential therapeutic implications for triple-negative breast cancer"

**A.**

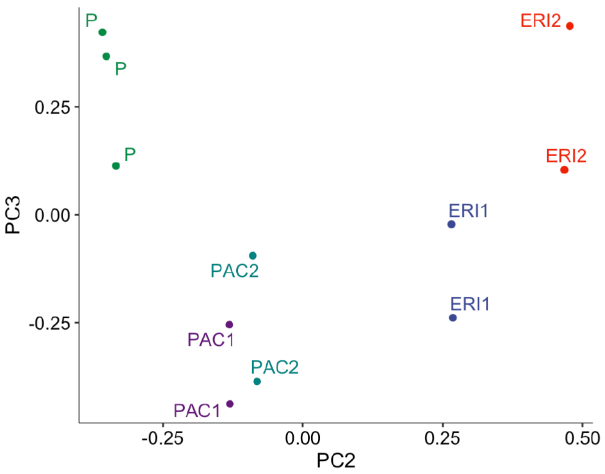


**Supplementary Figure 1. Global DNA methylation profiles cluster by treatment type.** PC1 and PC2 were determined by principal component analysis (PCA) on epigenome-wide methylation profile.

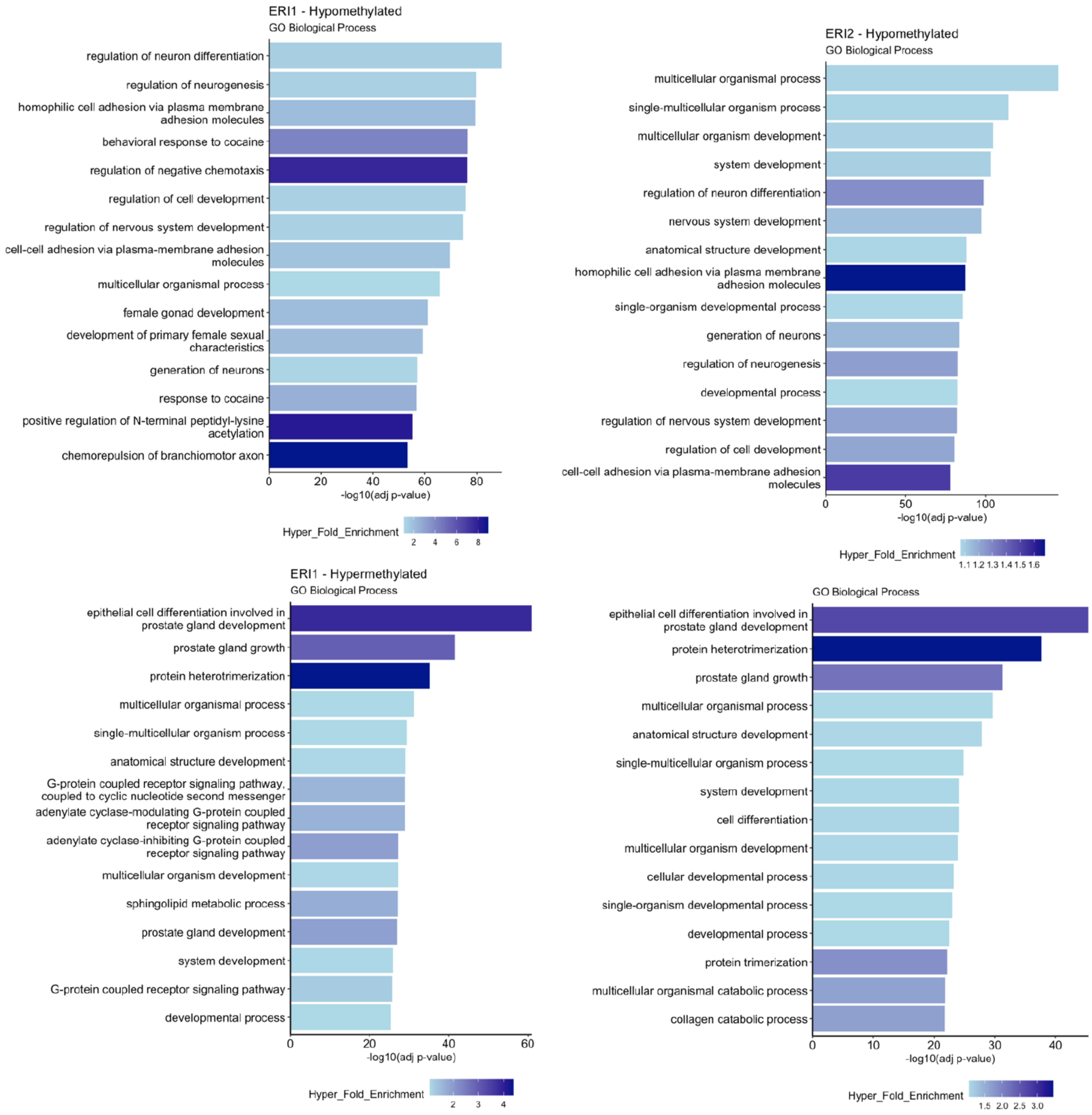


**Supplementary Figure 2. Additional eribulin treatment increases alterations in DNA methylation patterns.** Top 15 biological processes associated with genomic regions of differentially methylated CpGs associated with first eribulin treatment and second eribulin treatment as determined by Genomic Regions Enrichment of Annotations Tool (GREAT).

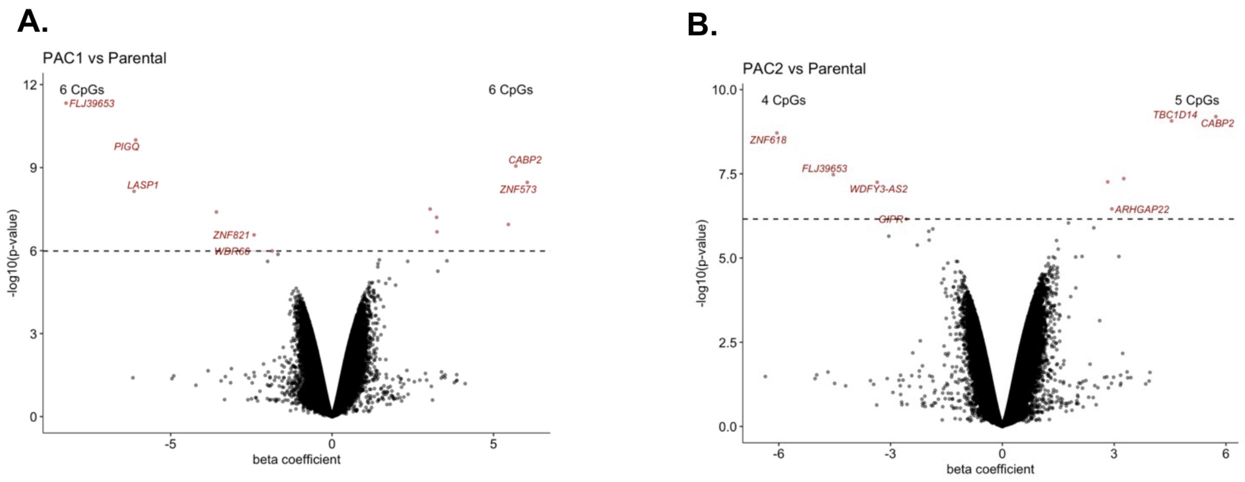


**Supplementary Figure 3. Eribulin treatment associated with differentially methylated CpGs are enriched in regulatory regions in DNA.** (A) Volcano plot of dmCpGs associated with first paclitaxel treatment compared to the methylation status of parental cells determined by an epigenome-wide association study. (B) Volcano plots of dmCpGs associated with second paclitaxel treatment compared to the methylation status of parental cells determined by an epigenome-wide association study. Colored in red are considered to be differentially methylated at q-value < 0.05.

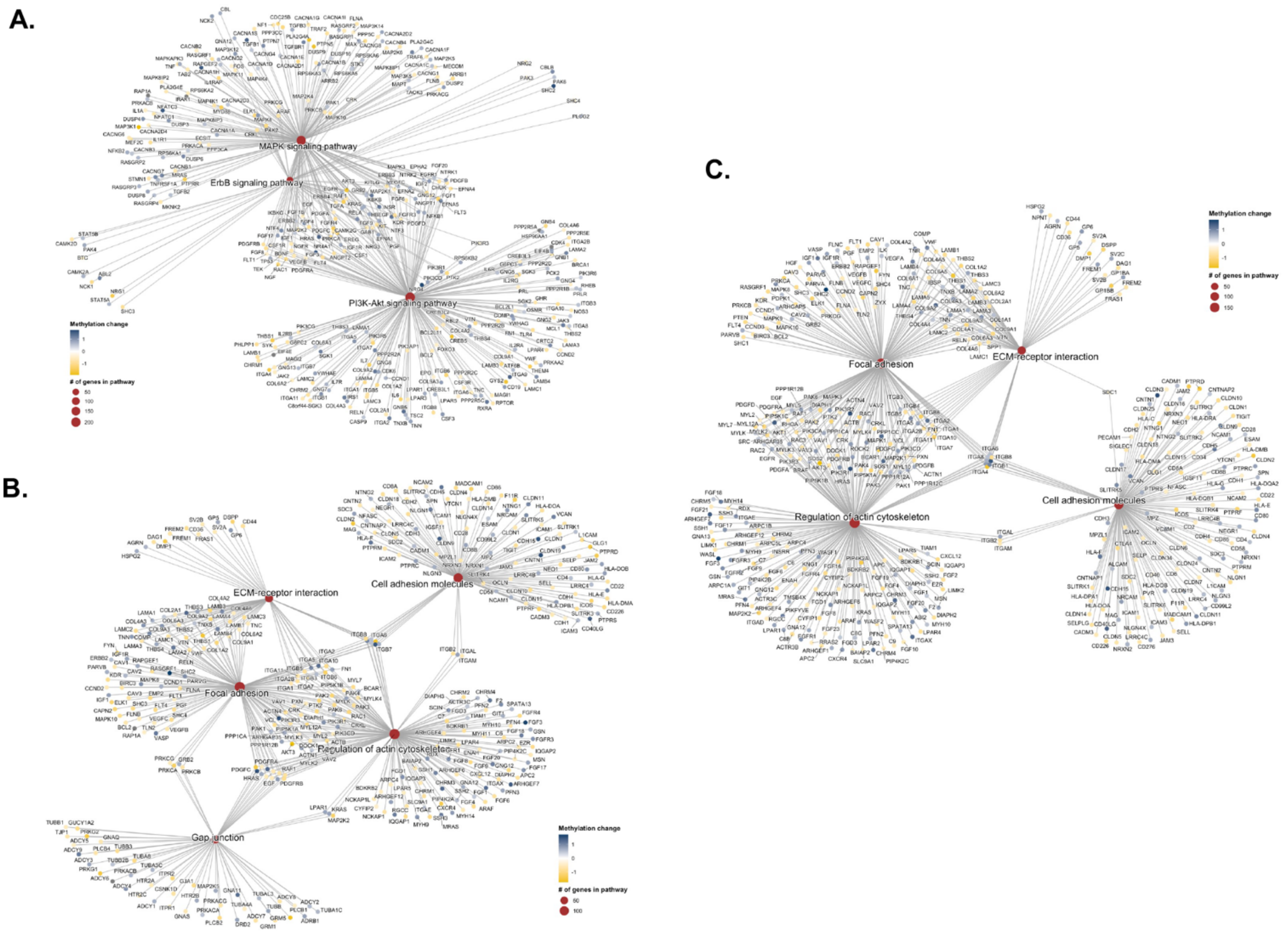


**Supplementary Figure 4. Genes with differentially methylated CpGs are associated with key biological processes in the mesenchymal-to-epithelial transition.** (A) ErbB-related signaling pathways were associated with genes containing dmCpGs from the first eribulin treatment. Extracellular matrix-related processes were significantly associated with genes containing differentially methylated CpGs after (B) the first eribulin treatment and (C) the second eribulin treatment.

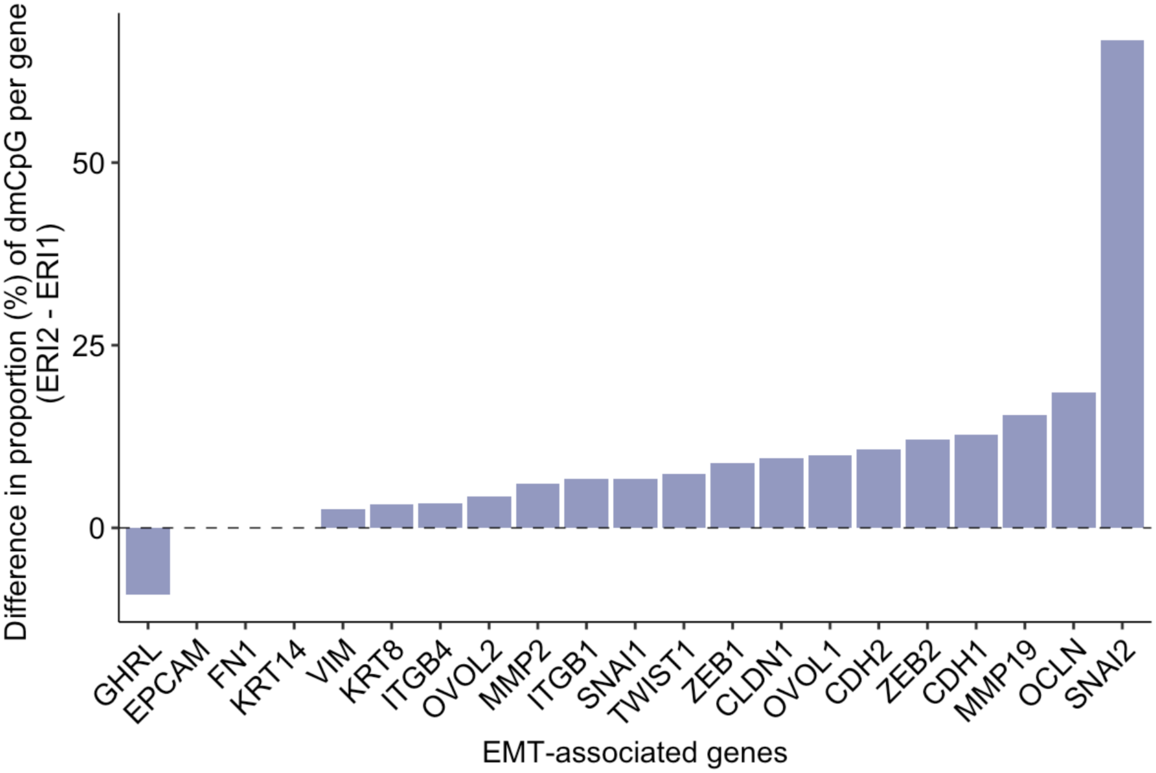


**Supplementary Figure 5. Differential methylation patterns are detected in Mesenchymal to Epithelial Transition-associated genes.** Change in proportion of dmCpGs in each of EMT-associated genes from the first eribulin treatment to the second eribulin treatment.

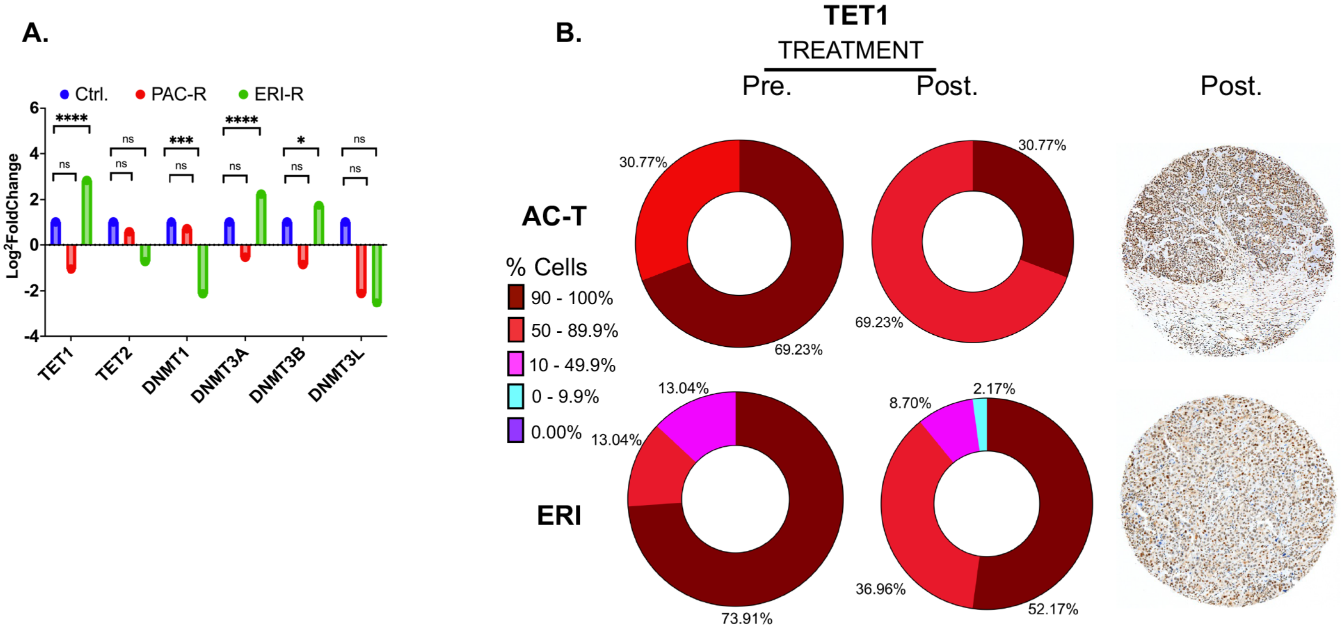


**Supplementary Figure 6. Eribulin treatment in primary human triple-negative breast cancers alters the expression of DNA methylation regulatory enzymes.** (A) Quantitative RT-PCR was performed to assess the expression levels of DNA methylation markers in the MDA MB-231 parental line and its resistant counterparts. (B) Representative images and quantification of TET1 were obtained through immunohistochemistry (IHC) analysis of specimens treated with AC-T and ERI.
